# Branch point control at malonyl-CoA node: A computational framework to uncover the design principles of an ideal genetic-metabolic switch

**DOI:** 10.1101/847947

**Authors:** Peng Xu

## Abstract

Living organism is an intelligent system encoded by hierarchically-organized information to perform precisely-controlled biological functions. Biophysical models are important tools to uncover the design rules underlying complex genetic-metabolic circuit interactions. Based on a previously engineered synthetic malonyl-CoA switch (Xu *et al*, PNAS 2014), we have formulated nine differential equations to unravel the design principles underlying an ideal metabolic switch to improve fatty acids production in *E. coli*. By interrogating the physiologically accessible parameter space, we have determined the optimal controller architecture to configure both the metabolic source pathway and metabolic sink pathway. We determined that low protein degradation rate, medium strength of metabolic inhibitory constant, high metabolic source pathway induction rate, strong binding affinity of the transcriptional activator toward the metabolic source pathway, weak binding affinity of the transcriptional repressor toward the metabolic sink pathway, and a strong cooperative interaction of transcriptional repressor toward metabolic sink pathway benefit the accumulation of the target molecule (fatty acids). The target molecule (fatty acid) production is increased from 50% to 10-folds upon application of the autonomous metabolic switch. With strong metabolic inhibitory constant, the system displays multiple steady states. Stable oscillation of metabolic intermediate is the driving force to allow the system deviate from its equilibrium state and permits bidirectional ON-OFF gene expression control, which autonomously compensates enzyme level for both the metabolic source and metabolic sink pathways. The computational framework may facilitate us to design and engineer predictable genetic-metabolic switches, quest for the optimal controller architecture of the metabolic source/sink pathways, as well as leverage autonomous oscillation as a powerful tool to engineer cell function.

## Introduction

In recent years, there is an influx of applying dynamic control theory to optimize metabolic pathways for production of various chemicals (Venayak, Anesiadis et al. 2015, Xu 2018, Xia, Ling et al. 2019). The marriage of intelligent control with synthetic biology have fruited a large volume of experimental and computational works that allow us to embrace a “dynamic” perspective to engineer cell metabolism (Zhang, Carothers et al. 2012, Xu, Li et al. 2014, Gupta, Reizman et al. 2017). The notion of “metabolic homeostasis” is a result of the dynamic interplay of the various biomolecules inside the cell (Xu 2018, Lv, Qian et al. 2019). Take the glycolytic pathway as an example, oscillating metabolic flux could arise due to the feedback inhibition of the phosphofructokinase by cellular energy levels (specifically, ATP, ADP and AMP) (Sel’kov 1968, Bier, Bakker et al. 2000, Chandra, Buzi et al. 2011, Gustavsson, van Niekerk et al. 2014). Another classical example is the Lac operon, hysteresis and multiple steady states could arise due to the positive feedback loop of the intake of the inducer (IPTG or lactose) by lactose permease encoded by *Lac*Y (Yildirim and Mackey 2003, Santillán, Mackey et al. 2007, Stamatakis and Mantzaris 2009). Inspired by this phenomena, early synthetic biology effort is spent extensively on constructing artificial genetic circuits by mimicking the electrical counterparts of the physical word (Andrianantoandro, Basu et al. 2006). Combing with mathematical modeling, a collection of classical work has emerged in the early 2000s, including the well-known toggle switch (CHEN and BAILEY 1994, Gardner, Cantor et al. 2000), repressilator (Elowitz and Leibler 2000) and metabolator (Fung, Wong et al. 2005) *et al*. These seminal works have encouraged us to employ biophysical models to quantitatively unravel and test the complicated molecular interactions underlying many perplexing biological problems, which marks the birth of synthetic biology.

With about one decade, the post-term impact of synthetic biology starts yielding fruits in the metabolic engineering field (Keasling 2010). From a control perspective, metabolic enzyme could be the “actuator” that performs chemical conversion (i.e. kinase phosphorylation, chromatin deacetylation) or the “transducer” that generates secondary messenger (i.e. cAMP or acetyl-CoA) (Smolke and Silver 2011, Michener, Thodey et al. 2012). Moving beyond the logic circuits engineering (AND, OR, NOT, NOR gates *et al*) (Tamsir, Tabor et al. 2011, Wang, Kitney et al. 2011, Moon, Lou et al. 2012), metabolic engineers have been able to harness various regulatory mechanisms, including repression (Liu, Xiao et al. 2015), activation (Doong, Gupta et al. 2018), attenuation (Benzinger and Khammash 2018) or RNA silencing (Yang, Lin et al. 2018), to rewire carbon flux and dynamically control cell metabolism. A number of control architectures (Oyarzún and Stan 2013, Liu, Xiao et al. 2015, Oyarzún and Chaves 2015, Venayak, Anesiadis et al. 2015, Chaves and Oyarzún 2019) have emerged and been applied to relieve metabolic burden (Ceroni, Boo et al. 2018), eliminate intermediate toxicity (Xu, Li et al. 2014), decouple cell growth from metabolite production (Bothfeld, Kapov et al. 2017, Doong, Gupta et al. 2018), eliminate metabolic heterogeneity (Xiao, Bowen et al. 2016, Rugbjerg, Myling-Petersen et al. 2018, Rugbjerg, Sarup-Lytzen et al. 2018, Wang and Dunlop 2019). The interdisciplinary connection among control theory, genetic principles, ecological and evolutional rules open a new venue for us to design and engineer precisely controlled genetic-metabolic circuits to reprogram biological functions (Calles, Goñi-Moreno et al. 2019). Engineering such decision-making functions to rewire the genetic (information) flow to redirect/optimize metabolic flux will enable us to deliver intelligent microbes for a broad range of applications, ranging from biocomputation, bioremediation, biosensing, biosynthesis to therapeutics (Nikel, Chavarría et al. 2016, Gao, Xu et al. 2019, Grozinger, Amos et al. 2019).

One of the essential tasks for metabolic engineers is to dynamically allocate carbon flux, so that the limited cellular resources could be harnessed to maximize the production of the target molecules (Xu, Bhan et al. 2013, Wan, Marsafari et al. 2019). Considering that the cell’s goal is to proliferate, there is always a tradeoff or conflicts between cell growth and metabolite overproduction. This will require us to equip the cells with various sensors to detect a broad range of environmental cues, cellular stimuli or metabolite intermediates (Zhang, Jensen et al. 2015, Wan, Marsafari et al. 2019), in such a way the cell can autonomously adjust gene expression or cell metabolism to compensate the loss or eliminate the surplus of enzyme activity. To achieve this, a number of control architectures, including the incoherent feedforward loop (Dunlop, Keasling et al. 2010, Harrison and Dunlop 2012), the invertor gate (Liu, Xiao et al. 2015), the metabolic toggle switch (Soma, Tsuruno et al. 2014) and the metabolic valve (Solomon and Prather 2011), have been implemented to improve the cellular tolerance to biofuels, or improve chemical production.

One of the highly studied dynamic control system is centering around the malonyl-CoA node (Xu, Li et al. 2014, Fehér, Libis et al. 2015, Albanesi and de Mendoza 2016, David, Nielsen et al. 2016). Malonyl-CoA is the essential metabolic building blocks for synthesizing advanced biofuels (Xu, Gu et al. 2013), lipids (Qiao, Wasylenko et al. 2017, Xu, Qiao et al. 2017), polyketides (Zhou, Qiao et al. 2010, Liu, Marsafari et al. 2019), oleochemicals (Xu, Qiao et al. 2016), flavonoids (Xiu, Jang et al. 2017) and cannabinoids (Luo, Reiter et al. 2019) *et al*. High level of malonyl-CoA benefits the production of these metabolites (Yang, Kim et al. 2018) but also inhibits cell growth (Xu, Li et al. 2014, Liu, Xiao et al. 2015). Up to date, the FapR-derived malonyl-CoA sensor has been successfully applied to mammalian cell (Ellis and Wolfgang 2012), *E. coli* (Xu, Wang et al. 2014, Yang, Kim et al. 2018) and yeast (Li, Si et al. 2015, David, Nielsen et al. 2016). In particular, a recent development of the malonyl-CoA oscillator (Xu, Li et al. 2014) has garnered significant attractions and allows us to study the optimal configurations of the controller architecture (**Fig. 1**). By integrating genetic and metabolic circuits, we have been able to experimentally construct and validate a malonyl-CoA oscillatory switch that was engineered to improve fatty acids production in *E. coli* (Xu, Li et al. 2014). Experimentally, we have engineered malonyl-CoA-responsive promoters that could be upregulated or down-regulated by FapR, and the activation or the repression could be reversed by malonyl-CoA. A recent study reported the same control structure with pGAP promoter and FapR regulator (Wen, Tian et al. 2020), and engineered a synthetic malonyl-CoA oscillator and metabolator in *Komagataella phaffii*. This dual direction ON-OFF control mimics the amino acid feedforward and feedback regulation that are naturally occurring in many bacteria.

**Fig. 1.**
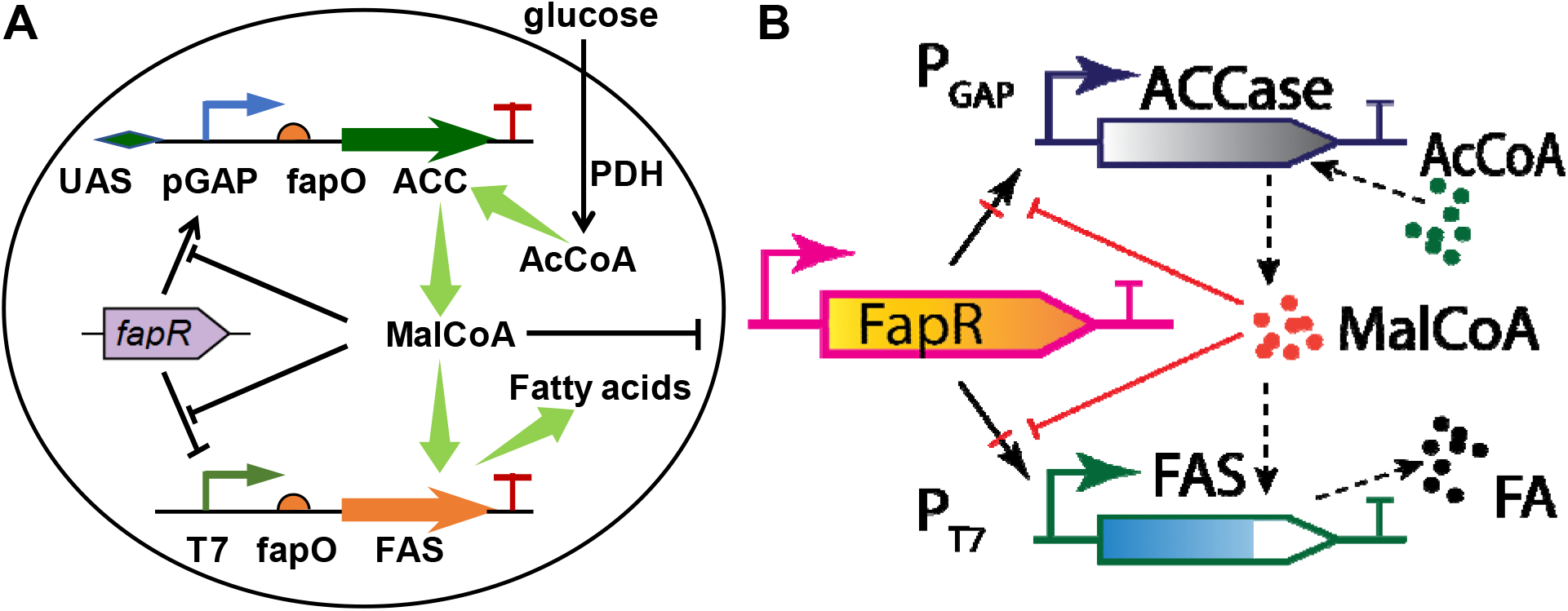
A malonyl-CoA switch to dynamically control fatty acids biosynthesis. (A) Autonomous ON-OFF control of malonyl-CoA. FapR activates pGAP promoter and upregulates the transcription of the malonyl-CoA source pathway (ACC) which generates malonyl-CoA; FapR represses T7 promoter and shuts down the transcription of the malonyl-CoA sink pathway (FAS) which consumes malonyl-CoA. The FapR bindings sites on the ACC operon is an upstream activation sequence (UAS). The FapR binding sites on the FAS operon is the fapO operator. Malonyl-CoA is the effector molecule (ligand) that antagonizes the activity of FapR. (B) Four possible genetic configurations of malonyl-CoA controller, which could be explored by changing the sign of the Hill coefficients (*n* and *p*) listed in Eqn. 4 and Eqn. 5. The black arrow with red cross indicates either transcriptional activation or repression.

One essential question is how to effectively configure the regulatory architecture of the metabolic source pathway and the metabolic sink pathway. To unravel the design principles underlying the malonyl-CoA switch, we set about to establish a biophysical model (system of ODE equations) and interrogated a broad range of parameter spaces, including the protein degradation rate (*D*), malonyl-CoA inhibitory constant (1/*K*_1_) and malonyl-CoA source pathway induction rate (*β*_2_). We also determined the optimal regulatory architecture for both the malonyl-CoA source pathway (ACCase) and the malonyl-CoA sink pathway (FAS), defined by the FapR-UAS dissociation constant (*K*_4_), FapR-fapO dissociation constant (*K*_3_) as well as the FapR-fapO Hill cooperativity coefficient (*n*). Our aim in this work is to understand how autonomous oscillation may contribute to optimal metabolite (fatty acids) production in strain engineering. The computational framework may facilitate us to design and engineer predictable genetic-metabolic switches, quest for the optimal controller architecture of the metabolic source/sink pathways, as well as leverage autonomous oscillation as a powerful tool to engineer cell function.

## Computational method and system equations

### Assumptions to develop the system equations

To simplify the biochemical and genetic events, we made eight assumptions to extract the basics of the genetic-metabolic circuits (**Fig. 1**): (***a***) We assume the number of DNA binding sites, specifically, FapO and UAS, far exceeds the number of transcriptional factor FapR in the system. Therefore, the repression rate of FAS or the activation rate of ACC are independent of the number of FapO and UAS in the system. (***b***) Glycolytic pathway (9 reactions) could be lumped into one single reaction to forming acetyl-CoA from glucose by PDH. (***c***) Fatty acids biosynthesis could be lumped into one single reaction to forming fatty acids (FA) from malonyl-CoA by FAS. (***d***) Malonyl-CoA depletion rate due to the formation of malonyl-CoA-FapR complex is negligible in the mass balance equation of malonyl-CoA (Eqn. 7). (***e***) The total enzyme or FapR concentration are approximately equivalent to the free enzyme or free FapR concentrations. (***f***) For non-regulated protein production (i.e. FapR and PDH), the production rate is cell growth-associated, therefore the production rate is proportional to the cell growth rate. (***g***) For regulated protein production (i.e. FAS and ACC), the production rate consists of both leaky expression (which is growth-associated) and regulated expression (which is non growth-associated) in the mass balance equations. (***h***) The cytosol is a homogenous and well-mixed system without mass transfer or diffusion limitations, where *D* could be interpreted as the dilution rate for CSTR or degradation constant for batch culture.

### Formulation of the kinetic rate and mass balance equations

We formulated the kinetic rate models (**Table 1**) on the basis of Michaelis-Mention equation for enzyme-substrate equations, Monod kinetics (Xu 2020) with metabolite (Malonyl-CoA) inhibition for cell growth, Hill-type equations for enzyme kinetics and metabolite-TF binding. Specifically, Eqn. 1 describes the specific growth rate, which follows Monod growth with glucose as limiting nutrients and malonyl-CoA as inhibitory factor; Eqn. 2 describes the mass balance for cell growth; Eqn. 3 describes the growth-associated production of FapR and the depletion of FapR due to the formation of FapR-Malonyl-CoA complex; Eqn. 4 describes the growth-associated production (leaky expression) of FAS and the regulated expression of FAS repressed by FapR; Eqn. 5 describes the growth-associated production (leaky expression) of ACC and the regulated expression of ACC activated by FapR; Eqn. 6 describes the production rate of fatty acids (FA) from malonyl-CoA; Eqn. 7 describes the mass balance for malonyl-CoA, accounting for both the malonyl-CoA source (ACC) pathway and the malonyl-CoA sink (FAS) pathway; Eqn. 8 describes the mass balance for acetyl-CoA, accounting for both the acetyl-CoA source (PDH) pathway and the acetyl-CoA sink (ACC) pathway; Eqn. 9 describes the PDH production rate which is proportional to the cell growth rate; and Eqn. 10 describes the mass balance for glucose, accounting for the consumption rate due to cell growth and acetyl-CoA production. For all the mass balance equations (Eqn. 2 to Eqn. 10), we also considered the dilution or degradation terms. Biomass and cell concentration in the feeding stream of the system were designated as *S*_0_ and *X*_0_.

**Table 1.** Equations used to define the autonomous oscillatory genetic-metabolic circuits.

| Equation No. | Equations used in this work |
| --- | --- |
| 1 | $\mu = \frac{\mu_{\max} S}{\left(\frac{C_{\text{MalCoA}}}{K_1} + 1\right) (K_S + S)}$ |
| 2 | $\frac{\partial}{\partial t} X(t) = D (X_0 - X(t)) + \mu X(t)$ |
| 3 | $\frac{\partial}{\partial t} C_{\text{FapR}}(t) = \alpha_1 \mu X(t) - D C_{\text{FapR}}(t) - \frac{k_1 C_{\text{FapR}}(t) C_{\text{MalCoA}}(t)^m}{K_2^m + C_{\text{MalCoA}}(t)^m}$ |
| 4 | $\frac{\partial}{\partial t} C_{\text{FAS}}(t) = \frac{\beta_1}{\left(\frac{C_{\text{FapR}}(t)}{K_3}\right)^n + 1} - D C_{\text{FAS}}(t) + \alpha_2 \mu X(t)$ |
| 5 | $\frac{\partial}{\partial t} C_{\text{ACC}}(t) = \frac{\beta_2}{\left(\frac{K_4}{C_{\text{FapR}}(t)}\right)^p + 1} - D C_{\text{ACC}}(t) + \alpha_3 \mu X(t)$ |
| 6 | $\frac{\partial}{\partial t} C_{\text{FA}}(t) = \frac{k_2 C_{\text{FAS}}(t) C_{\text{MalCoA}}(t)^q}{K_m^q + C_{\text{MalCoA}}(t)^q} - D C_{\text{FA}}(t)$ |
| 7 | $\frac{\partial}{\partial t} C_{\text{MalCoA}}(t) = \frac{k_3 C_{\text{ACC}}(t) C_{\text{AcCoA}}(t)^r}{K_5^r + C_{\text{AcCoA}}(t)^r} - D C_{\text{MalCoA}}(t) - \frac{k_2 C_{\text{FAS}}(t) C_{\text{MalCoA}}(t)^q}{Y_{\text{PS1}} (K_m^q + C_{\text{MalCoA}}(t)^q)}$ |
| 8 | $\frac{\partial}{\partial t} C_{\text{AcCoA}}(t) = \frac{k_4 C_{\text{PDH}}(t) S(t)^u}{K_6^u + S(t)^u} - \frac{k_3 C_{\text{ACC}}(t) C_{\text{AcCoA}}(t)^r}{K_5^r + C_{\text{AcCoA}}(t)^r} - D C_{\text{AcCoA}}(t)$ |
| 9 | $\frac{\partial}{\partial t} C_{\text{PDH}}(t) = \alpha_4 \mu X(t) - D C_{\text{PDH}}(t)$ |
| 10 | $\frac{\partial}{\partial t} S(t) = D (S_0 - S(t)) - \frac{\mu X(t)}{Y_{XS}} - \frac{k_4 C_{PDH}(t) S(t)^u}{Y_{PS2} (K_6^u + S(t)^u)}$ |

### Computational methods

Matlab R2017b was used as the computational package on a Windows 7 professional operation system. The CPU processor is Intel Core i3-6100 with 3.70 GHz. The installed memory (RAM) is 4.0 GHz. Matlab symbolic language package coupled with LaTex makeup language were used to compile the equations (Table 1). ODE45 solver was used to simulate and predict the system behavior. Matlab plot function was used to output the solutions and graphs. Matlab codes will be shared upon request. Biological parameters for Fig. 2 to Fig. 10 could be found in the supplementary files. Most of the parameters were assigned on the basis of BioNumbers database (Milo, Jorgensen et al. 2009). Jacobian matrices were evaluated according to a reported numerical method (Auralius Manurung (2020). Calculate Jacobian of a function numerically at a given condition (https://www.github.com/auralius/numerical-jacobian)).

**Fig. 2.**
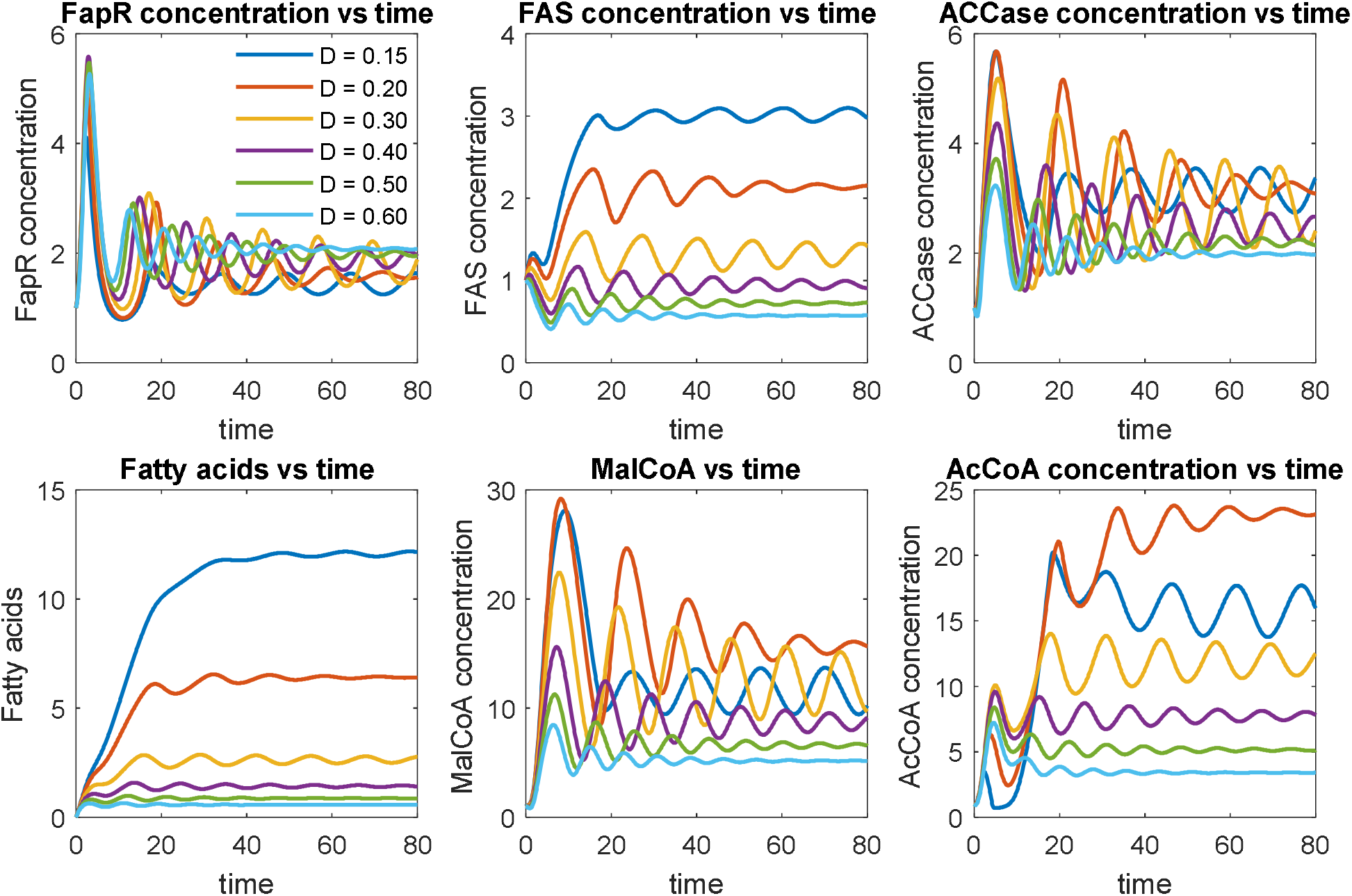
Effect of protein/metabolite degradation rate (dilution rate) on system dynamic behavior. Protein degradation rates have been labelled with different color. Low degradation rate (*D* = 0.15) leads to relatively stable oscillation. High degradation rate (*D* ≥ 0.2) leads to damped oscillation. The units are arbitrary units. Parameters could be found in the supplementary file.

Initial states determine the final states for dynamic system. In this work, the initial conditions were taken on the basis of physiologically accessible dataset of biochemical systems. Most of the numbers were consistent with biochemical engineering textbooks, including Shuler & Kargi, Bioprocess engineering; and Blanch & Clark, Biochemical Engineering *et al*. These initial conditions come with SI unit and is provided in the SI file.

## Results and Discussion

### Effect of protein/metabolite degradation rate (dilution rate) on system dynamic behavior

To understand the system dynamics, we probed a number of parameter space to generate the dynamic pattern that meets our design and control criteria. A list of parameters could be found in the supplementary files. We first investigated how protein/metabolite degradation rate impacts the system dynamics (**Fig. 2**). For all the simulations, we used six species, including regulator protein FapR, fatty acid synthase (FAS), acetyl-CoA carboxylase (ACCase), target product fatty acids (FA), intermediates malonyl-CoA (MalCoA) and acetyl-CoA (AcCoA), to represent the system.

Under the prescribed parameter conditions (supplementary files) with protein degradation rates ranging from 0.15 to 0.60 (the unit is inverse of time), we evaluated the trajectory of the numerical solutions of the system ODE equations (**Table 1**). For relatively high degradation rate (*D* ≥ 0.2), we observed that the system solutions are approximately behaving like a damped oscillator (**Fig. 2**). On the other hand, the low degradation rate (or longer residence time, i.e. *D* = 0.15 in **Fig. 2**) allows the system to oscillate stably with fixed frequency and amplitude, leading to the highest fatty acids production (**Fig. 2**). For example, fatty acids production at low protein degradation rate (*D* = 0.15) is about 10-folds higher than the fatty acids production at high protein degradation rate (*D* = 0.6). This is not counterintuitive as low degradation rate allows the protein catalysts stay longer in the system (Gao, Hou et al. 2019). And the stable oscillation indicates that the designed control scheme could perform alternating ON-OFF control of the malonyl-CoA source pathway and malonyl-CoA sink pathway. Interestingly, the fatty acids production pattern is closely related with the malonyl-CoA sink pathway (FAS), but doesn’t correlate well with the activity of the malonyl-CoA source pathway (ACC). This is rooted in our initial assumptions that sufficient malonyl-CoA will inhibit cell growth. As a result, the intermediate acetyl-CoA and malonyl-CoA displays distinct oscillating pattern, with the stable oscillation (*D* = 0.15) leading to better control.

We also explored whether we could further improve fatty acids production by using even smaller degradation rate (*i.e. D* = 0.1, **Fig. 3**). Interestingly, decreasing the degradation rate to 0.1 allows FapR to quickly accumulate in the system from *t* = 20. We could notice that a spike of fatty acids production at *t* = 20, but the entire control system collapses (*D* = 0.1, **Fig. 3**) at *t* > 20, due to the overdosed FapR repressing the expression of the malonyl-CoA sink pathway (FAS). Accompanying with increased FapR, the malonyl-CoA source pathway (ACCase) was also overdosed (*D* = 0.1, **Fig. 3**) due to the activating action of FapR toward the expression of ACCase. However, malonyl-CoA was not accumulated in the system due to the antagonist effect of FapR toward malonyl-CoA. Taken together, the low degradation rate (*D* = 0.1) allows the cell to only build biomass, but generates little final products (Fatty acids in this study). In summary, the range of degradation rate of the sensor protein (FapR) and the malonyl-CoA source pathway (ACCase) determines whether the designed control scheme will work or fail.

**Fig. 3.**
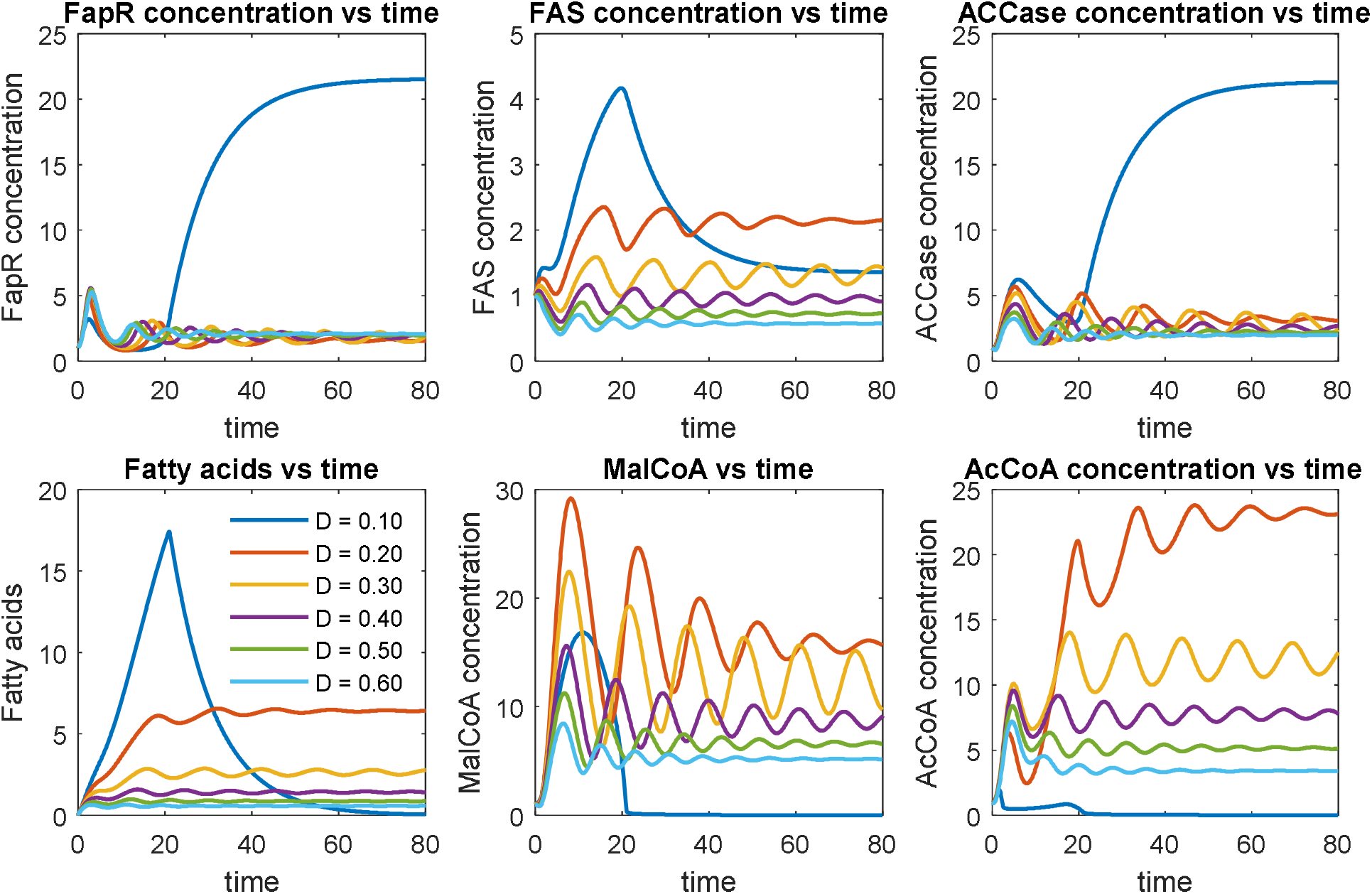
Effect of protein/metabolite degradation rate on system dynamic behavior. Protein degradation rates have been labelled with different color. Low degradation rate (*D* = 0.10) leads to a collapsed system: too much FapR represses the expression of FAS, activates the expression of ACC and quickly antagonize the resulting malonyl-CoA at *t* > 20.

Phase-plane represents the solution constraints between the interacting components, at different parameter conditions (such as dilution rate or binding affinity) (Xu 2020). We further performed a phase-plane analysis to interrogate the solutions of above ODEs (**Fig. 4**). On the FAS-FapR phase plane, the system is attracted to periodic limit cycle of clockwise motion. The horizontal (x-axis) projection of the elliptic cycle forms a negative slope with FapR (x-axis), indicating that FapR represses the expression of FAS. On the ACCase-FapR phase planes (**Fig. 4**), the system is attracted to periodic limit cycle of counterclockwise motion. The horizontal (x-axis) projection of the elliptic cycle forms a positive slope with FapR, indicating that FapR activates the expression of ACCase. Similarly, on the MalCoA-FapR phase plane (**Fig. 4**), the system is attracted to periodic limit cycle of counterclockwise motion. The horizontal (x-axis) projection of the elliptic cycle forms a negative slope with FapR, indicating that FapR acts as an antagonist for malonyl-CoA. Under *D* = 0.15, we observed that the system leads to stable oscillation (Fig. S1).

**Fig. 4.**
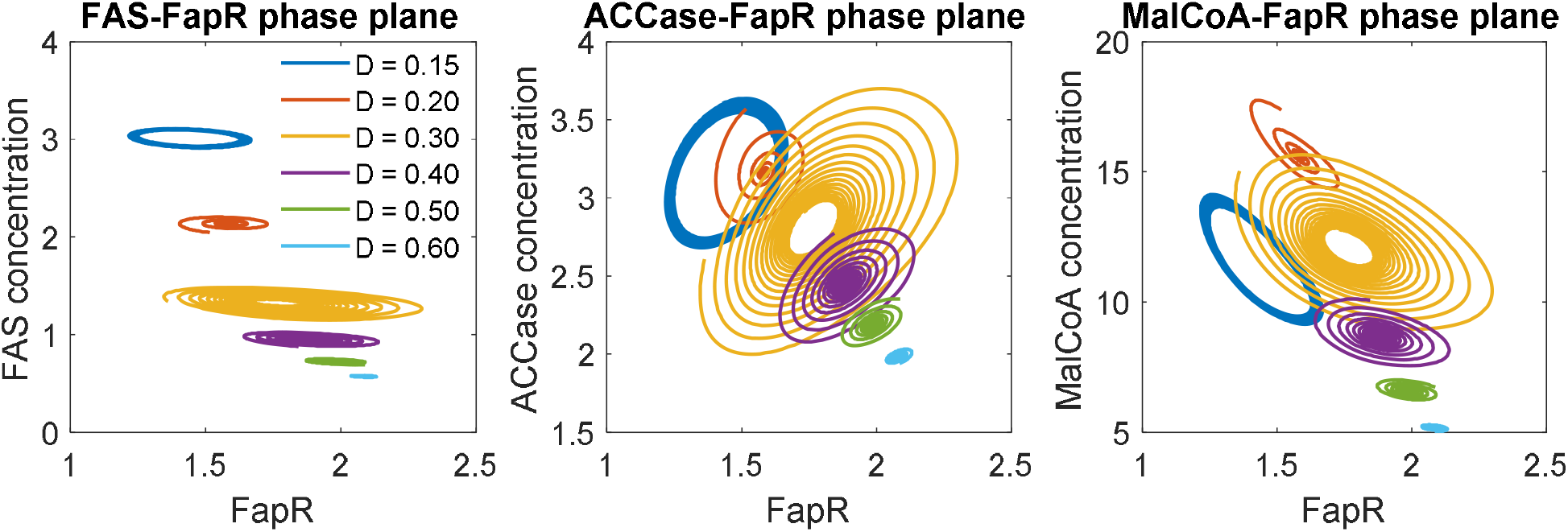
The phase-plane correlations for FAS-FapR, ACCase-FapR and MalCoA-FapR. FAS-FapR phase plane shows periodic limit cycle of clockwise motion. ACCase-FapR and MalCoA-FapR phase planes show periodic limit cycle of counterclockwise motions.

To analytically identify the steady state, we need to derive the Jacobina matrix and analyze the eigenvalue of the Jacobian matrix at each of the steady states. If all the eigenvalues are negative or the real parts of all eigenvalues are negative (for imaginary eigenvalues), this will be a stable steady state. Graphically, steady states represent time-invariant solutions along the time-axis. The trajectory of stable steady states will asymptotically or periodically converge to a fixed point or travel on an orbit (a limit cycle). By analyzing the Jacobian matrices, two pure imaginary eigenvalues with zero real parts were arrived (Supplementary Notes 1), indicating a stable oscillation under *D* = 0.15. The phase portraits allow us to understand the motion of system dynamic behavior, it may also serve as diagnosis for troubleshooting the design-build-test cycle in genetic circuit engineering.

### Effect of malonyl-CoA dissociation constant (*K*_1_) on system dynamic behavior

We next investigated how the malonyl-CoA dissociation constant (*K*_1_) impacts the system dynamics (**Fig. 5**). The malonyl-CoA dissociation constant (*K*_1_) describes the inhibitory strength of malonyl-CoA to cell growth: small dissociation constant (*K*_1_) indicates a high binding affinity and high inhibitory strength. A number of dissociation constants ranging from 0.10 to 4.0 (in the units of concentration) were investigated (**Fig. 5**). As expected, strong inhibition (*K*_1_ = 0.10) will sequestrate the cell at a low growth rate and lead to constant expression of FapR, FAS and ACCase (**Fig. 5**), indicating that the expression of FAS and ACCase are independent of the control scheme. As the inhibition becomes weaker (*K*_1_ = 0.50 and 1.00), the solution of the system ODEs oscillates with increased amplitude, albeit the frequency of the oscillation remains unchanged. A perfect ON and OFF control of FAS and ACCase expression is taking place when a medium strength of inhibition (*K*_1_ = 2.0) is used. This medium strength of inhibition confers the system to oscillate stably with improved fatty acids production (**Fig. 5**), albeit the fatty acids increase is less than 50%. When the dissociation constant takes a larger number (*K*_1_ = 4.0), the systems behave like a damped oscillation that is approximately approaching to the optimal design scheme (*K*_1_ = 2.0). This analysis indicates that a medium strength of dissociation constant (*K*_1_) should be used. In practice, one can always use adaptive lab evolution to screen conditionally tolerant phenotype that meets the *K*_1_ selection criteria.

**Fig. 5.**
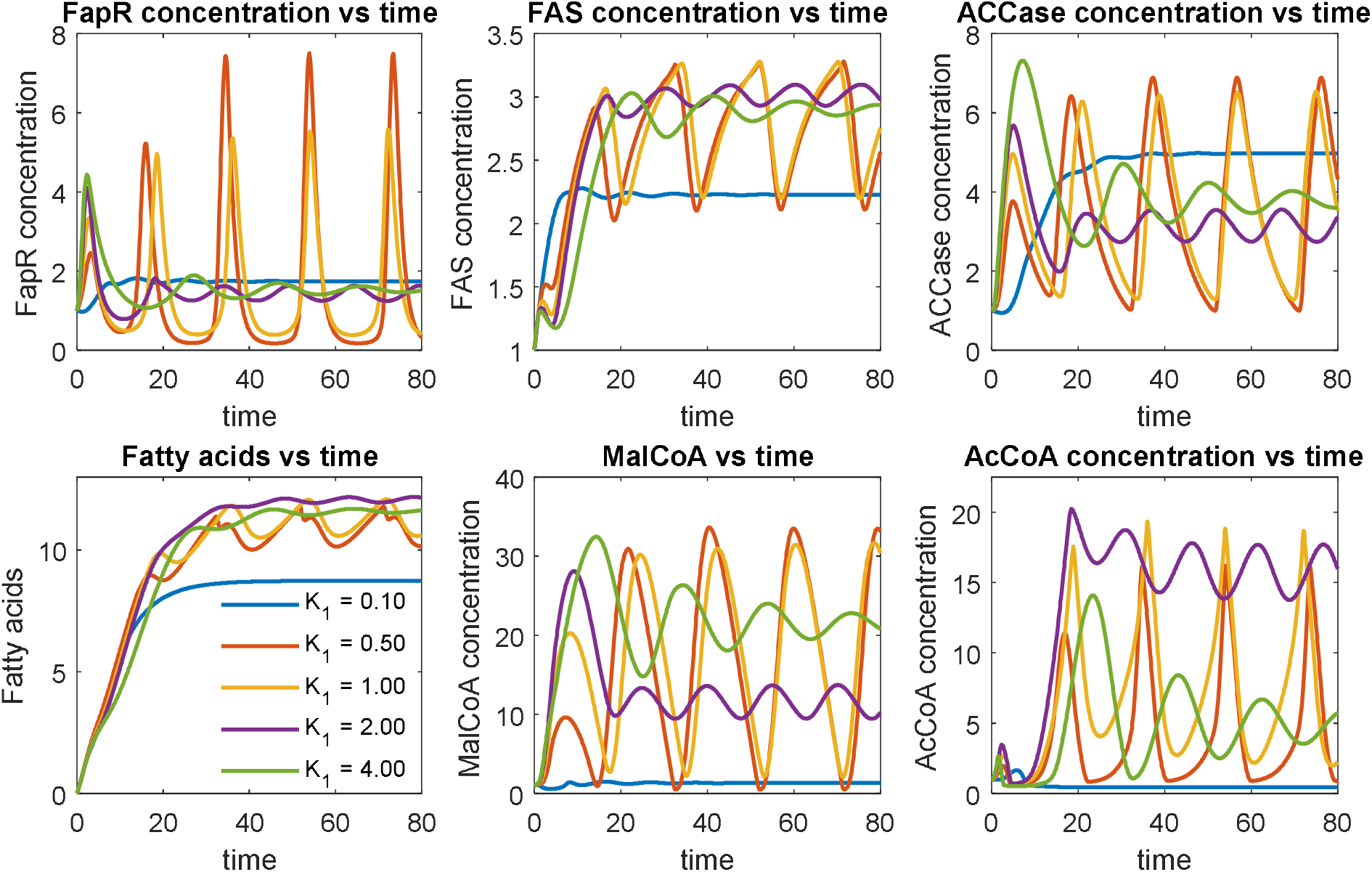
Effect of malonyl-CoA dissociation constant (*K*_1_) on system dynamic behavior. Malonyl-CoA dissociation constants (*K*_1_) have been labelled with different color. Low dissociation constants (*K*_1_ = 0.5, 1.0 and 2.0) lead to stable oscillation. High dissociation constant (*K*_1_ = 4.0) leads to damped oscillation. Medium strength of malonyl-CoA inhibition (*K*_1_ = 2.0) favors fatty acids production.

Similarly, we could perform a phase-plane analysis (**Fig. 6**). The phase-planes suggest that the optimal control scheme (*K*_1_ = 2.0, the purple cycles) only permits a very narrowed space of FAS, ACCase and MaloCoA solutions. Interestingly, for low malonyl-CoA dissociation constant (*K*_1_= 0.5), the system exhibits a looping behavior on the FAS-FA and MalCoA-FA phase plane. Plotting the steady state solutions of fatty acids, FAS and malonyl-CoA, we observed looping pattern of solutions in the 3-D space, this may also imply a hysteretic state of the system (Aris, Borhani et al. 2019) (Supplementary Notes 2 and 3). It simply means that strong malonyl-CoA inhibition (i.e. *K*_1_= 0.3 or 0.5) will lead to multiplicity of steady states (**Fig. 6 and Fig. 7**), which is a critical factor to evaluate the dynamics of the system behavior.

**Fig. 6.**
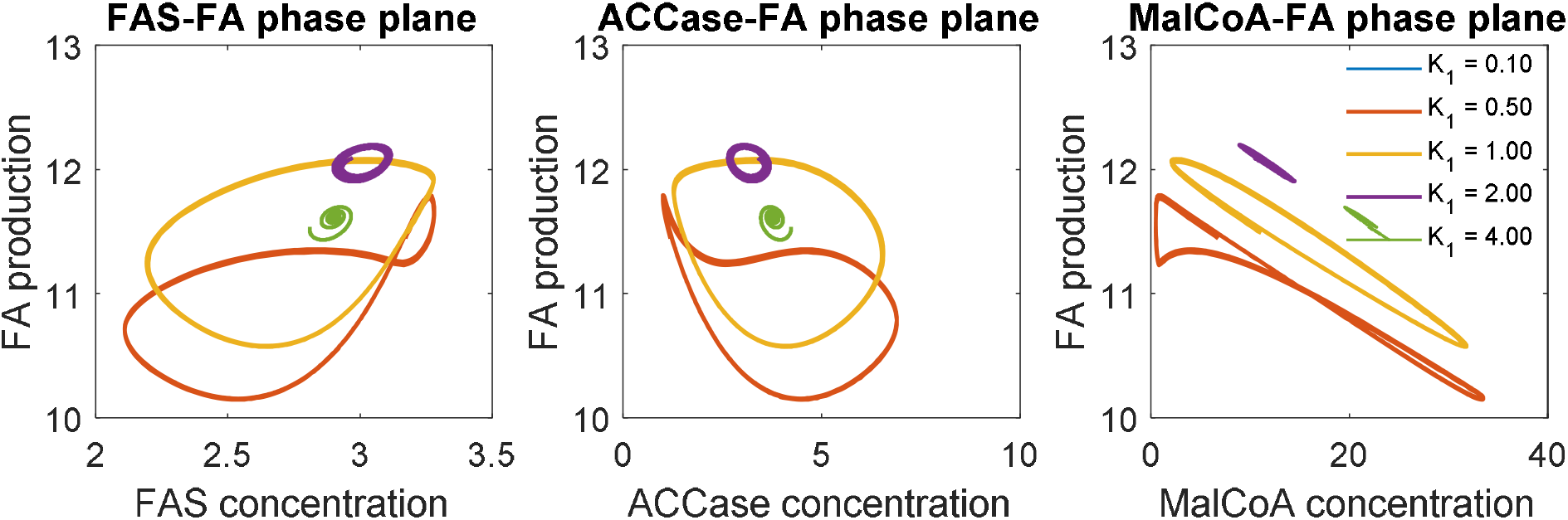
The phase-plane portraits for FA-FAS, FA-ACCase and FA-MalCoA. Low malonyl-CoA dissociation constant (*K*_1_= 0.5, orange line), which corresponds to strong malonyl-CoA inhibition, leads to multiplicity of steady states pattern between FAS-FA and MalCoA-FA input-output relationships.

**Fig. 7.**
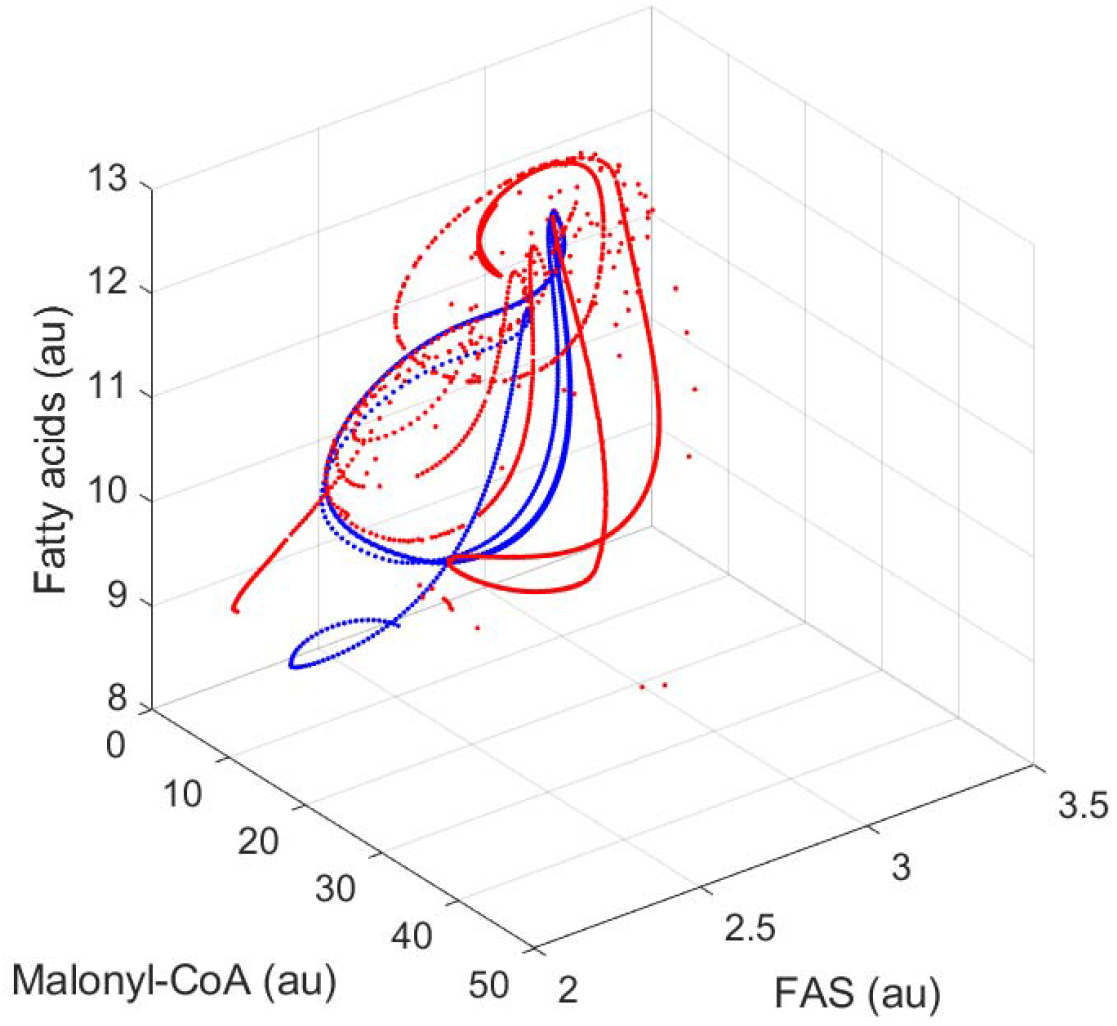
The 3-D phase-plane portraits for fatty acids, FAS and malonyl-CoA, with malonyl-CoA inhibition constant (*K*_1_) varying from 0.1 to 1.5. A specific trajectory for *K*_1_ = 0.3 is added to the above solution space from a random point (1,1,1), marked in blue color. Other parameters used here are the same as the parameters used in Fig. 5.

Literature reports that feedback inhibition of free fatty acid on FAS complex plays a major role in regulating FA synthesis. Specifically, it is generally believed that acyl-ACPs or acyl-CoAs will feedback inhibit acetyl-CoA carboxylase in *E. coli* (Davis and Cronan 2001). Since acyl-CoA/ACP could be hydrolyzed to free fatty acids by acyl-CoA thioesterase *tes*A (which is constitutively overexpressed in the published paper PNAS 2014), we believe the feedback inhibition of acyl-CoA/ACP on FAS complex could be minimized when *tes*A was overexpressed. In this synthetic system, the malonyl-CoA inhibitory effect on FAS was translated to the malonyl-CoA inhibitory effect on cell growth: cell growth is associated with how much of membrane lipids (phospholipids synthesized from acyl-CoAs/acyl-ACPs) were made. Therefore, the malonyl-CoA/ACP feedback inhibitory effect on FAS (cell growth) plays a critical role to determine the system dynamics.

### Effects of FapR-UAS interaction on system dynamics

We next explored how the gene expression of the malonyl-CoA source pathway (ACCase) impacts the system dynamics. According to the original design and Eqn.5, expression of ACCase is governed by the FapR-UAS interactions. The system equation for ACCase (Eqn. 5) account for both the growth-associated leaky expression (*α*_3_) and the FapR-activated regulatory expression (*β*_2_, *p* and *K*_4_). In all our simulations, we assume stringent regulation and the leaky expression is negligible (*α*_2_ = *α*_3_ = 0.05). We will specifically investigate how the ACCase induction rate (*β*_2_) and the FapR-UAS dissociation constant (*K*_4_) impact the system dynamic (**Fig. 8** and **Fig. 9**).

**Fig. 8.**
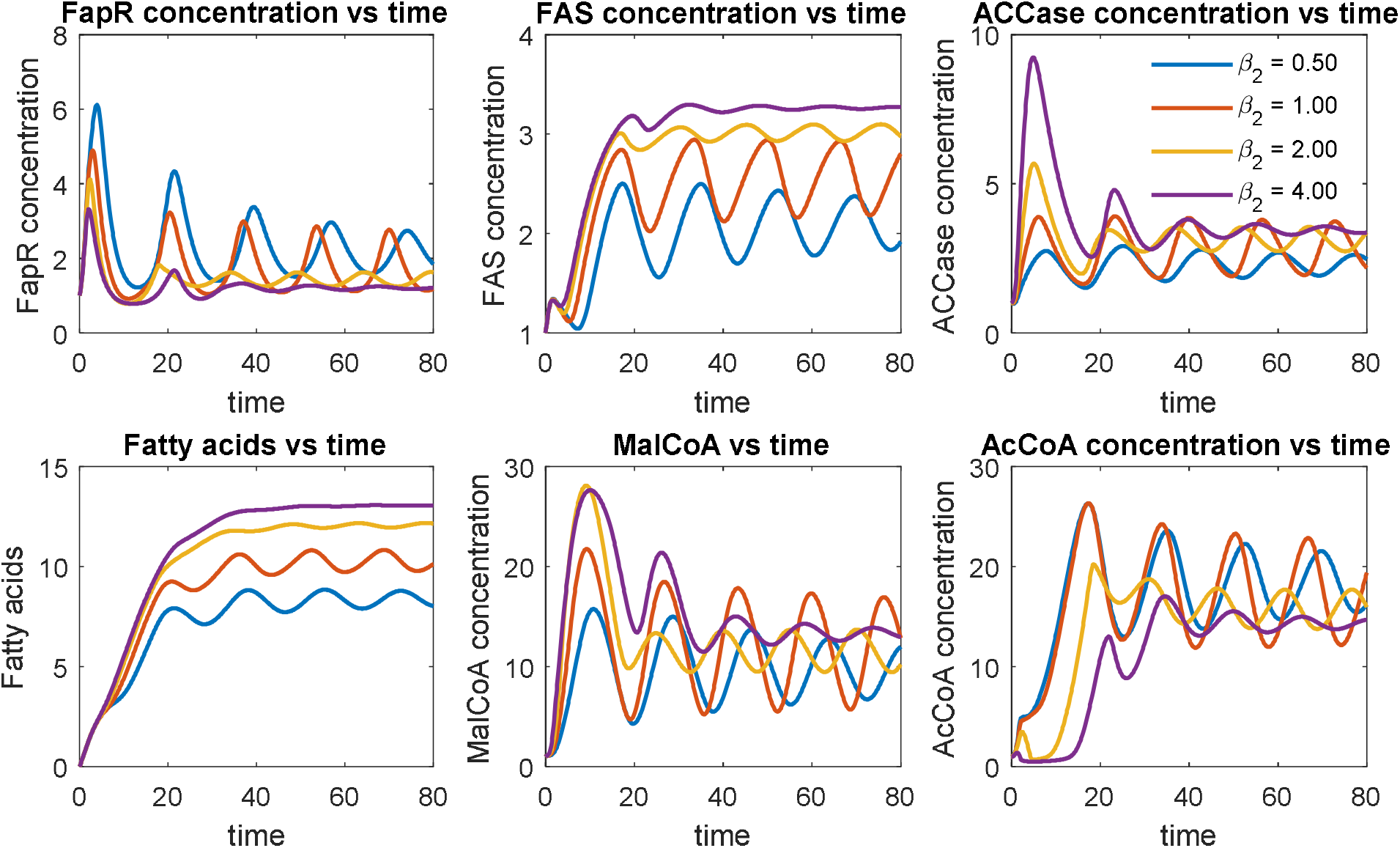
Effect of ACCase induction rate (*β*_2_) on system dynamics. ACCase induction rates (*β*_2_) have been labelled with different color. High ACCase induction rate (i.e. *β*_2_ = 4.0) leads to a quickly damped oscillation and favors fatty acid accumulation.

**Fig. 9.**
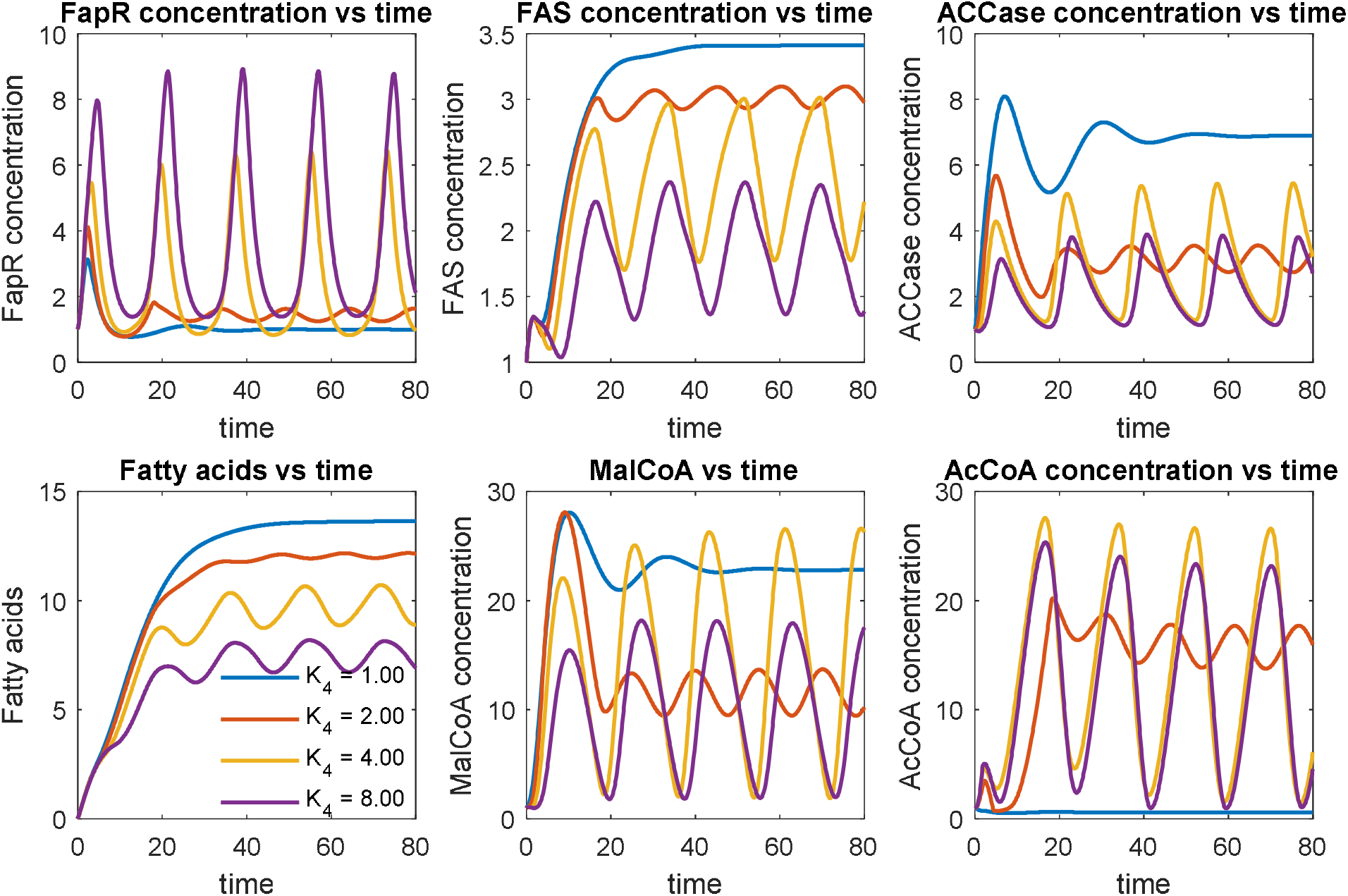
Effect of FapR-UAS dissociation constant (*K*_4_) on system dynamics. Tighter FapR-UAS binding is advantageous to fatty acids production.

We investigated a number of ACCase induction rates (*β*_2_, in the units of concentration per time), ranging from 0.50 to 4.0 (**Fig. 8**). As the ACCase induction rate increases (*β*_2_) from 0.50 to 4.0, the expression of malonyl-CoA source pathway (ACCase) is upregulated, leading to improved fatty acids production (**Fig. 8**). For example, the fatty acids production is increased up to 2-fold when the ACCase induction rate (*β*_2_) increases from 0.5 (blue line, **Fig. 8**) to 4.0 (purple line, **Fig. 8**). On the other hand, the amount of regulator protein FapR decreases with increasing ACCase induction rate (*β*_2_) (**Fig. 8**), possibly due to the antagonist effect of malonyl-CoA. However, this monotonic correlation was not found for the species MalCoA and AcCoA, due to the complicated autoregulation of malonyl-CoA in the control system. Furthermore, under low ACCase induction rates (i.e. *β*_2_ = 0.5, 1.0 and 2.0), the oscillation damped periodically with decreasing amplitude. Under high ACCase induction rate (i.e. *β*_2_ = 4.0), the oscillation damped quickly to reach its steady state (**Fig. 8**). This result indicates that a high ACCase induction rate (*β*_2_) is essential for the proper function of the control scheme.

As FapR is the activator for the ACC operon, and the DNA binding site for FapR is a UAS (upstream activation sequence). We next investigated how the FapR-UAS dissociation constant (*K*_4_) impacts the system dynamics (**Fig. 9**). A smaller FapR-UAS dissociation constant (*K*_4_) indicates a tighter binding between FapR and UAS (the inverse of the dissociation constant quantifies the binding affinity). As the binding between FapR and UAS becomes tighter (*K*_4_ decreases from 8.0 to 1.0), the expression of the malonyl-CoA source pathway (ACCase) is strongly activated, leading to increased fatty acids production (**Fig. 9**). For example, the fatty acids production is increased up to 2.2-fold when the FapR-UAS dissociation constant (*K*_4_) decreases from 8.0 (purple line, **Fig. 9**) to 1.0 (blue line, **Fig. 9**). Under high FapR-UAS binding affinity (*K*_4_ = 1.0), the oscillation damped quickly to reach its steady state; under low FapR-UAS binding affinity (*K*_4_ = 4.0 or 8.0), the oscillation retains periodic pattern with fixed frequency and amplitude. This result indicates that a tighter FapR-UAS binding is the critical factor to achieve the desired control scheme.

### Effect of FapR-fapO interaction on system dynamics

We also attempted to understand how the gene expression of the malonyl-CoA sink pathway (FAS) impacts the system dynamics. By design, FapR is the repressor that is specifically bound to fapO and represses the expression of the malonyl-CoA sink pathway (FAS). The system equation for FAS (Eqn. 4) accounts for both the growth-associated leaky expression (*α*_2_) and the FapR-repressed regulatory expression (*β*_1_, *n* and *K*_3_). Transcriptional factor (FapR) and DNA binding site (fapO) interactions are typically defined by the binding affinity (inverse of the dissociation constant) and the Hill cooperativity coefficient. By probing the physiologically accessible parameter space, we will investigate how the FapR-fapO dissociation constant (*K*_3_) impact the system dynamics (**Fig. 10**).

**Fig. 10.**
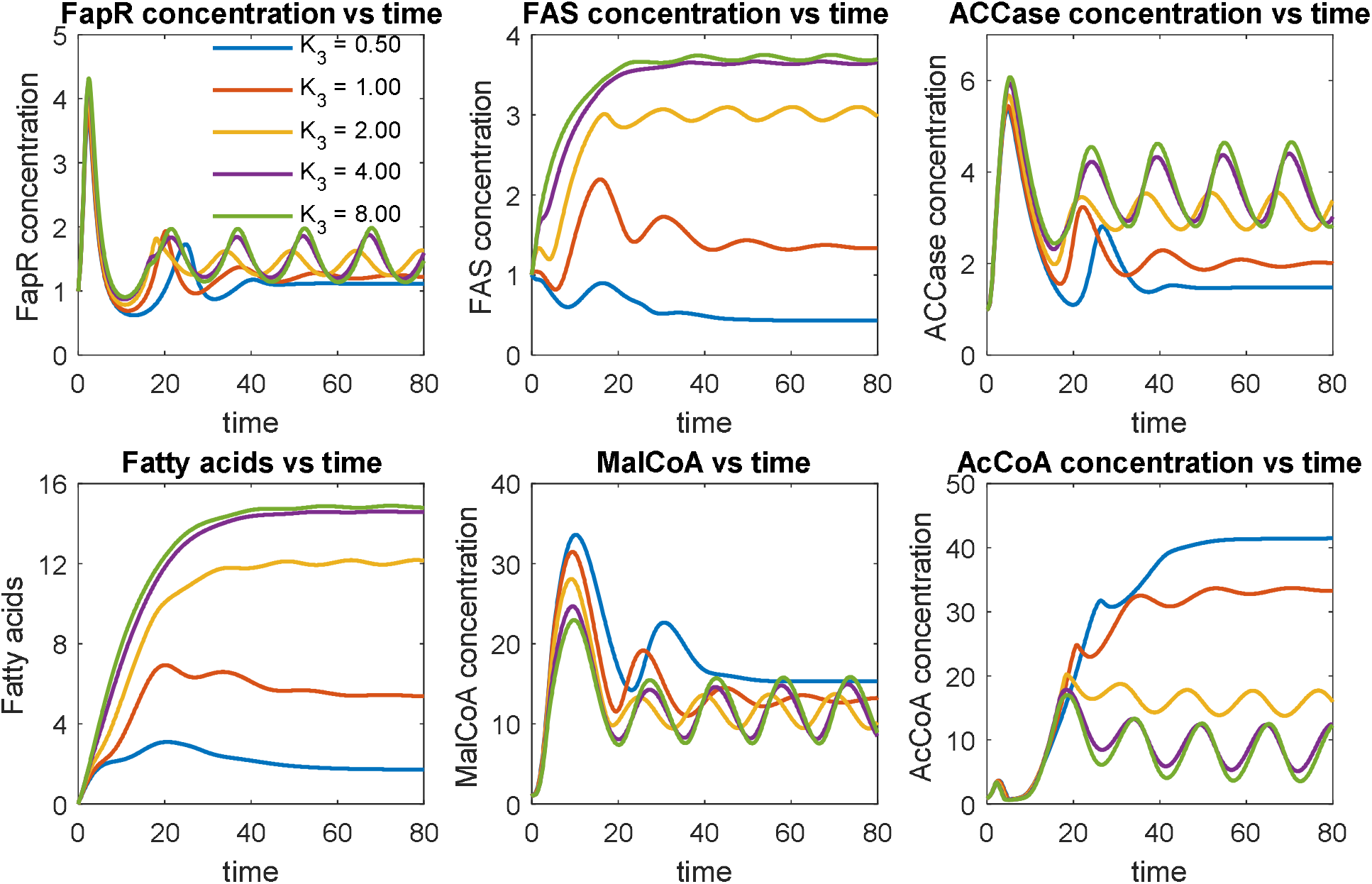
Effect of FapR-fapO dissociation constant (*K*_3_) on system dynamics. A weak binding between FapR and fapO (or a large FapR-fapO dissociation constant, i.e. *K*_3_ = 8.0) significantly improves fatty acid production, up to 7-fold.

We investigated a number of FapR-fapO dissociation constant (*K*_3_), spanning from 0.50 to 8.0 (**Fig. 10**). A smaller FapR-fapO dissociation constant indicates a tighter binding between FapR and fapO, thus the FapR-fapO complex will function as a stronger roadblock to prevent FAS transcription. As the binding between FapR and fapO becomes tighter (*K*_3_ decreases from 8.0 to 0.5), the expression of the malonyl-CoA sink pathway (FAS) is strongly repressed (**Fig. 10**), leading to decreased fatty acid accumulation. For example, the fatty acids production at low FapR-fapO dissociation constant (*K*_3_ = 0.50, blue curve) is less than 1/7 of the fatty acid production at high FapR-fapO dissociation constant (*K*_3_ = 8.0, green curve) (**Fig. 10**). With weaker FapR-fapO binding (*K*_3_ = 4.0 and 8.0), the ODE solutions for ACCase, MalCoA and AcCoA oscillate with fixed frequency and amplitude, indicating the functionality of the ON-OFF control toward both the malonyl-CoA source pathway (ACCase) and the malonyl-CoA sink pathway (FAS). However, with tighter FapR-fapO binding (*K*_3_ = 0.5 and 1.0), the oscillation collapses at relatively short period of time, indicating a faulted control scheme. This result suggests that a weak binding between FapR and fapO (or a large FapR-fapO dissociation constant) is the most important design criteria to achieve the desired ON-OFF control scheme.

### Exploring the optimal controller architecture

The Hill cooperativity coefficient is a critical factor determining the input-output relationship of biological signal transduction. Recent studies demonstrate that lots of nonlinear and complicated biological functions are arising from the cooperative assembly of biological molecules (Bashor, Patel et al. 2019, Shaw, Yamauchi et al. 2019), including DNA, RNA and proteins. As such, we will investigate how the FapR-FapO Hill cooperativity coefficient (*n*) impacts the system dynamics (**Fig. 11**). We choose a number of FapR-fapO Hill cooperativity coefficients, ranging from −4 to 4. It should be noted that, our original mass balance equations (Eqn. 4 and Eqn. 5) only account for the fact that FapR represses the expression of FAS and FapR activates the expression of ACC, which corresponds to a positive Hill coefficient (*n* > 0 and *p* > 0).

**Fig. 11.**
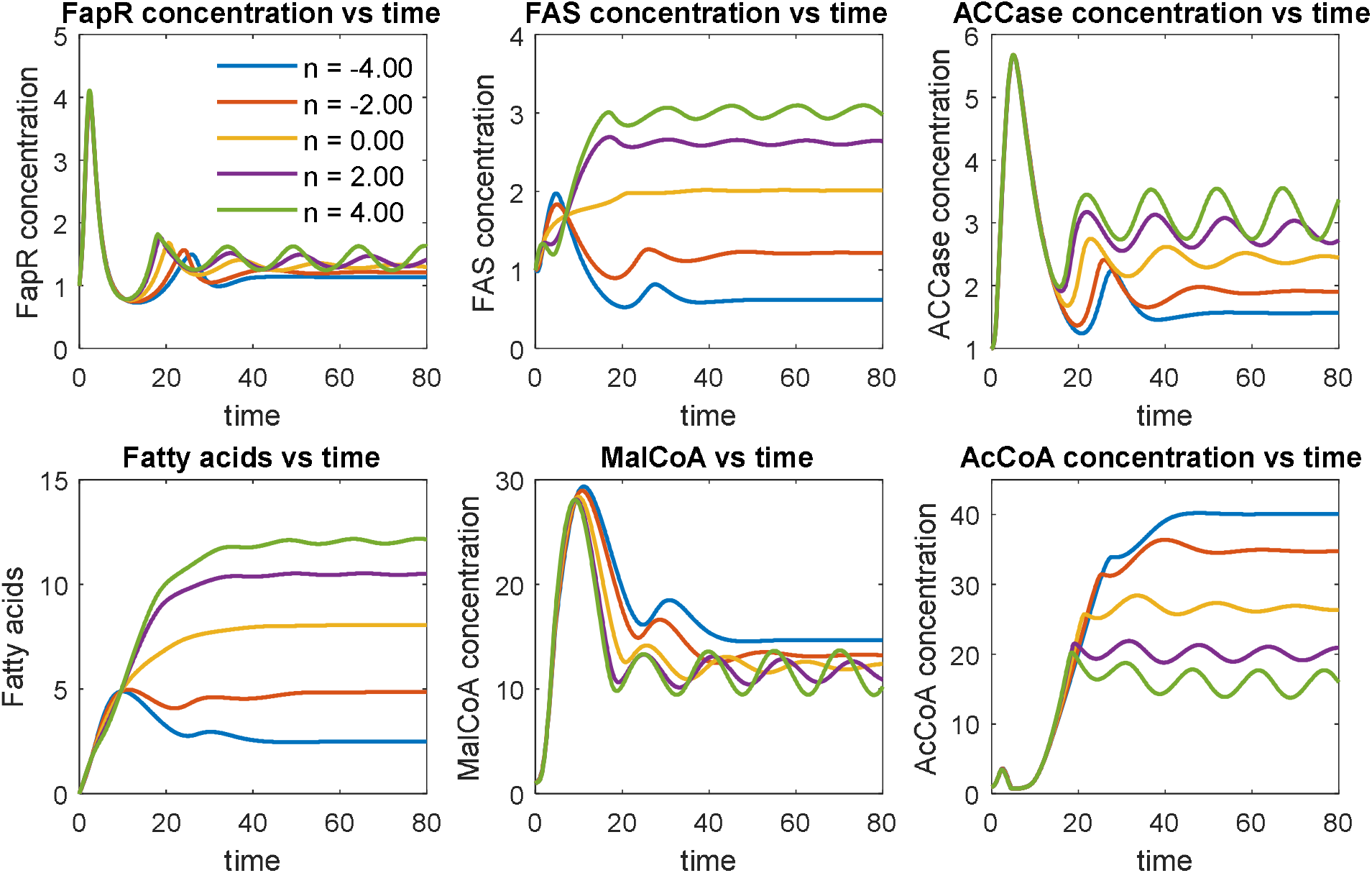
Effect of FapR-fapO Hill cooperativity coefficient (*n*) on system dynamics. Strong repression (*n* = 4) leads to stable oscillation and drives the cell to make more fatty acids.

The sign of the Hill coefficient is related with the genetic configuration of the controller (**Fig. 1B**). For example, a positive Hill coefficient (*n*) in the malonyl-CoA sink pathway (FAS) indicates that FapR *<u>represses</u>* the transcriptional activity of FAS expression; while a negative Hill coefficient (*n*) in the malonyl-CoA sink pathway (FAS) indicates that FapR *<u>activates</u>* the transcriptional activity of FAS expression (Equation 4). Similarly, a positive Hill coefficient (*p*) in the malonyl-CoA source pathway (ACC) indicates that FapR *<u>activates</u>* the transcriptional activity of ACC expression; while a negative Hill coefficient (*p*) in the malonyl-CoA source pathway (ACC) indicates that FapR *<u>represses</u>* the transcriptional activity of ACC expression (Equation 5). By changing the sign of the Hill coefficients for the malonyl-CoA sink pathway (FAS) and the malonyl-CoA source pathway (ACC), we could explore the ‘optimal controller’ structure in this study (**Fig. 1B**). Here we consider both positive Hill coefficients (*n* = 2.0 and 4.0) and negative Hill coefficients (*n* = −2.0 and −4.0) as well as no cooperation (*n* = 0). By comparing either the activating or repressing effect of FapR, we may interrogate the topology of the optimal controller architecture that leads to maximal fatty acids production.

As the FapR-fapO Hill cooperativity coefficient (*n*) increases from −4.0 to 4.0, the regulatory action of FapR towards the malonyl-CoA sink pathway (FAS) shifts from activation to repression. As a result, a significant increase in the FAS, ACCase expression and fatty acids production are observed (**Fig. 11**). For example, almost 5-fold increase of fatty acids is obtained when the FapR-fapO Hill cooperativity coefficient (*n*) increases from −4.0 (blue line, strong activation) to 4.0 (green line, strong repression). Under strong FapR activation (*n* = −4.0), counterintuitively, the expression of FAS is instead downregulated (**Fig. 11**). This could be linked to the unbalanced induction rate (*β*) between the malonyl-CoA source pathway (ACCase, *β*_2_ = 2.0) and the malonyl-CoA sink pathway (FAS, *β*_1_ = 0.5). Even with highly cooperative activation of FAS by FapR (*n* = −4.0, blue line), the low induction rate of the malonyl-CoA sink pathway (FAS) makes the expression of FAS unable to catch up with the expression of ACCase (malonyl-CoA source pathway). As a result, malonyl-CoA will build up but stay unchanged in the system (blue line in **Fig. 11**) to inhibit cell growth, which will result in even lower level of FapR (activator for FAS expression when *n* = −4, blue line) and therefore exacerbate the expression of FAS. On the contrary, highly cooperative repression of FAS by FapR (*n* = 4, green line) will make malonyl-CoA level oscillate, which forms the driving force to dynamically link and control the expression of the malonyl-CoA source pathway (ACCase) and the malonyl-CoA sink pathway (FAS). This analysis indicates that a control architecture consisting of upregulated metabolic source and downregulated metabolic sink is an essential design criterion to build adaptive genetic-metabolic circuits. In addition, the stable oscillation of the metabolic intermediate (i.e., malonyl-CoA) forms the driving force to exert the ON-OFF dynamic control toward complex metabolic function in the cell.

## Conclusions

With the better understanding of cellular regulation, metabolic engineers have been able to engineer both the chemistry (the mass flow) and the control modules (the information flow) inside the cell to design intelligent cell factories with improved performance. Moving beyond thermodynamic and stoichiometric constraints, living organisms could be viewed as a smart system consisting of sensor (ligand binding domain of transcriptional factors), transducers (DNA-binding domain of transcriptional factors, kinase or enzyme *et al*) and actuators (RNA polymerases). Along this direction, cellular regulation and feedback control mechanisms have been exploited to construct genetic/metabolic circuits that could sense/respond to environment, achieve adaptive metabolic function and reshape cell fate for diverse biotechnological and medical applications. As chemical engineers have done to program machine language and control the mass and energy flow in a chemical plant, a synthetic biologist could rewrite the genetic software and encode logic functions in living cells to control cellular activity.

Biophysical and biochemical models are important tools to quantitatively understand genetic circuit dynamics, metabolic network constraints, cell-cell communications (Dai, Lee et al. 2019) and microbial consortia interactions (Kong, Meldgin et al. 2018, Tsoi, Wu et al. 2018). Based on a previously engineered malonyl-CoA switch, nine differential equations were formulated (Table 1) and employed to unravel the design principles underlying a perfect metabolite switch. While the models present in the current study were simple, they provide sufficient kinetic information to predict the dynamic behavior of the published work. By interrogating the physiologically accessible parameter space, we have determined the optimal control architecture to configure both the malonyl-CoA source pathway and the malonyl-CoA sink pathway. We also investigated a number of biological parameters that strongly impact the system dynamics, including the protein degradation rate (*D*), malonyl-CoA inhibitory constant (1/*K*_1_), malonyl-CoA source pathway induction rate (*β*_2_), FapR-UAS dissociation constant (*K*_4_), FapR-fapO dissociation constant (*K*_3_) as well as the FapR-fapO Hill cooperativity coefficient (*n*). We identified that low protein degradation rate (*D*), medium strength of malonyl-CoA inhibitory constant (1/*K*_1_), high malonyl-CoA source pathway induction rate (*β*_2_), strong FapR-UAS binding affinity (1/*K*_4_), weak FapR-fapO binding affinity (1/*K*_3_) and a strong cooperative repression of malonyl-CoA sink pathway (FAS) by FapR (*n*) benefits the accumulation of the target molecule (fatty acids). The fatty acids production could be increased from 50% to 10-folds with the different set of parameters. Under certain conditions (i.e. strong malonyl-CoA inhibitory constant 1/*K*_1_), the system will display multiplicity of steady states. Stable oscillation of malonyl-CoA is the driving force to make the system perform the ON-OFF control and automatically adjust the expression of both the malonyl-CoA source (ACCase) and malonyl-CoA sink (FAS) pathways.

In this work, we have chosen a number of biophysical parameters to discuss the possible output of the malonyl-CoA switch. Genetically, these parameters could be altered by web-lab experiments, including protein engineering or degenerated repressor binding sites to change the biding affinity between the interacting components *et al*. The computational framework present here may facilitate us to design and engineer predictable genetic-metabolic switches, configure the optimal controller architecture of the metabolic source/sink pathways, as well as reprogram metabolic function for various applications.

## Supporting information

Supplementary Figures and Matlab code

## Acknowledgments

Dr. Xu would like to acknowledge the Cellular & Biochem Engineering Program of the National Science Foundation under grant no.1805139 for funding support. Dr. Xu would also like to acknowledge the discussion of this project with the ENCH482/682 and ENCH640 students at the University of Maryland Baltimore County in the Fall 2018 and Spring 2019.

## Appendix Symbols and variables used in this work

*μ*: specific growth rate
*μ*_max_: maximum specific growth rate
*α*_1_: cell growth-associated FapR production rate constant (constitutive expression)
*α*_2_: cell growth-associated FAS production rate constant (leaky expression)
*α*_3_: cell growth-associated ACC production rate constant (leaky expression)
*α*_4_: cell growth-associated PDH production rate constant (constitutive expression)
*β*_1_: non cell growth-associated FAS production rate (regulated expression)
*β*_2_: non cell growth-associated ACC production rate (regulated expression)
*K*_1_: Malonyl-CoA inhibitory (dissociation) constant
*K*_2_: Mal-CoA and FapR saturation constant
*K*_3_: dissociation rate constant of free FapR toward fapO in the FAS operon (to repress FAS transcription)
*K*_4_: dissociation rate constant of free FapR toward UAS in the ACC operon (to activate ACC transcription)
*K*_5_: acetyl-CoA saturation (Michaelis) constant toward ACC
*K*_6_: glucose saturation (Michaelis) constant toward glycolytic pathway
*K*_*S*_: Monod constant for glucose
*K*_*m*_: Malonyl-CoA saturation (Michaelis) constant toward FAS
*k*_1_: FapR-inactivating rate constant due to the formation of MalCoA-FapR complex
*k*_2_: FA (fatty acids) production rate constant from Mal-CoA catalyzed by FAS
*k*_3_: malonyl-CoA production rate constant from acetyl-CoA catalzyed by ACC
*k*_4_: acetyl-CoA production rate constant from glycolysis catalzyed by PDH
*S*: glucose concentration
*S*_0_: glucose concentration in the feeding stream
*D*: dilution rate or degradation rate
*X*_0_: biomass concentration in the feeding stream
*Y*_PS1_: malonyl-CoA to fatty acids conversion yield
*Y*_XS_: glucose to biomass conversion yield
*Y*_PS2_: glucose to acetyl-CoA conversion yield
*m*: malonyl-CoA-FapR (ligand-TF) Hill cooperativity coefficient
*n*: FapR-FapO nucleoprotein complex Hill cooperativity coefficient
*p*: FapR-UAS nucleoprotein complex Hill cooperativity coefficient
*q*: malonyl-CoA-FAS (substrate-enzyme) Hill cooperativity coefficient
*r*: acetyl-CoA-ACC (substrate-enzyme) Hill cooperativity coefficient
*u*: glucose-PDH (substrate-enzyme, artificial reaction) Hill cooperativity coefficient

