## Supplementary Figures and Matlab code for "Branch point control at malonyl-CoA node: A computational framework to uncover the design principles of an ideal genetic-metabolic switch"

Peng Xu*

Department of Chemical, Biochemical and Environmental Engineering, University of Maryland Baltimore County, Baltimore, MD 21250

**Parameters used for Fig. 1 to Fig. 10**

*Parameters for Fig. 2, Fig. 3 and Fig. 4*: alpha1=0.8; alpha2=0.05; alpha3=0.05; alpha4=0.8; beta1=0.5; beta2=2.0; m=4; n=4; p=4; q=4; r=4; u=4; K1=2; K2=5; K3=2; K4=2; K5=0.5; K6=0.5; k1=0.5; k2=0.6; k3=2; k4=2; X0=0; S0=45; Y_XS=0.6; Y_PS1=0.4; Y_PS2=1.8; mu_max=2.2; K_S=0.75; K_m=0.5;

*Parameters for Fig. 5 and Fig. 6*: alpha1=0.8; alpha2=0.05; alpha3=0.05; alpha4=0.8; beta1=0.5; beta2=2.0; m=4; n=4; p=4; q=4; r=4; u=4; D=0.15; K2=5; K3=2; K4=2; K5=0.5; K6=0.5; k1=0.5; k2=0.6; k3=2; k4=2; X0=0; S0=45; Y_XS=0.6; Y_PS1=0.4; Y_PS2=1.8; mu_max=2.2; K_S=0.75; K_m=0.5;

*Parameters for Fig. 7*: alpha1=0.8; alpha2=0.05; alpha3=0.05; alpha4=0.8; beta1=0.5; m=4; n=4; p=4; q=4; r=4; u=4; D=0.15; K1=2; K2=5; K3=2; K4=2; K5=0.5; K6=0.5; k1=0.5; k2=0.6; k3=2; k4=2; X0=0; S0=45; Y_XS=0.6; Y_PS1=0.4; Y_PS2=1.8; mu_max=2.2; K_S=0.75; K_m=0.5;

*Parameters for Fig. 8*: alpha1=0.8; alpha2=0.05; alpha3=0.05; alpha4=0.8; beta1=0.5; beta2=2.0; m=4; n=4; p=4; q=4; r=4; u=4; D=0.15; K1=2; K2=5; K3=2; K5=0.5; K6=0.5; k1=0.5; k2=0.6; k3=2; k4=2; X0=0; S0=45; Y_XS=0.6; Y_PS1=0.4; Y_PS2=1.8; mu_max=2.2; K_S=0.75; K_m=0.5;

*Parameters for Fig. 9*: alpha1=0.8; alpha2=0.05; alpha3=0.05; alpha4=0.8; beta1=0.5; beta2=2.0; m=4; n=4; p=4; q=4; r=4; u=4; D=0.15; K1=2; K2=5; K4=2; K5=0.5; K6=0.5; k1=0.5; k2=0.6; k3=2; k4=2; X0=0; S0=45; Y_XS=0.6; Y_PS1=0.4; Y_PS2=1.8; mu_max=2.2; K_S=0.75; K_m=0.5;

*Parameters for Fig. 10*: alpha1=0.8; alpha2=0.05; alpha3=0.05; alpha4=0.8; beta1=0.5; beta2=2.0; m=4; p=4; q=4; r=4; u=4; D=0.15; K1=2; K2=5; K3=2; K4=2; K5=0.5; K6=0.5; k1=0.5; k2=0.6; k3=2; k4=2; X0=0; S0=45; Y_XS=0.6; Y_PS1=0.4; Y_PS2=1.8; mu_max=2.2; K_S=0.75; K_m=0.5;

**Supplementary Figures**

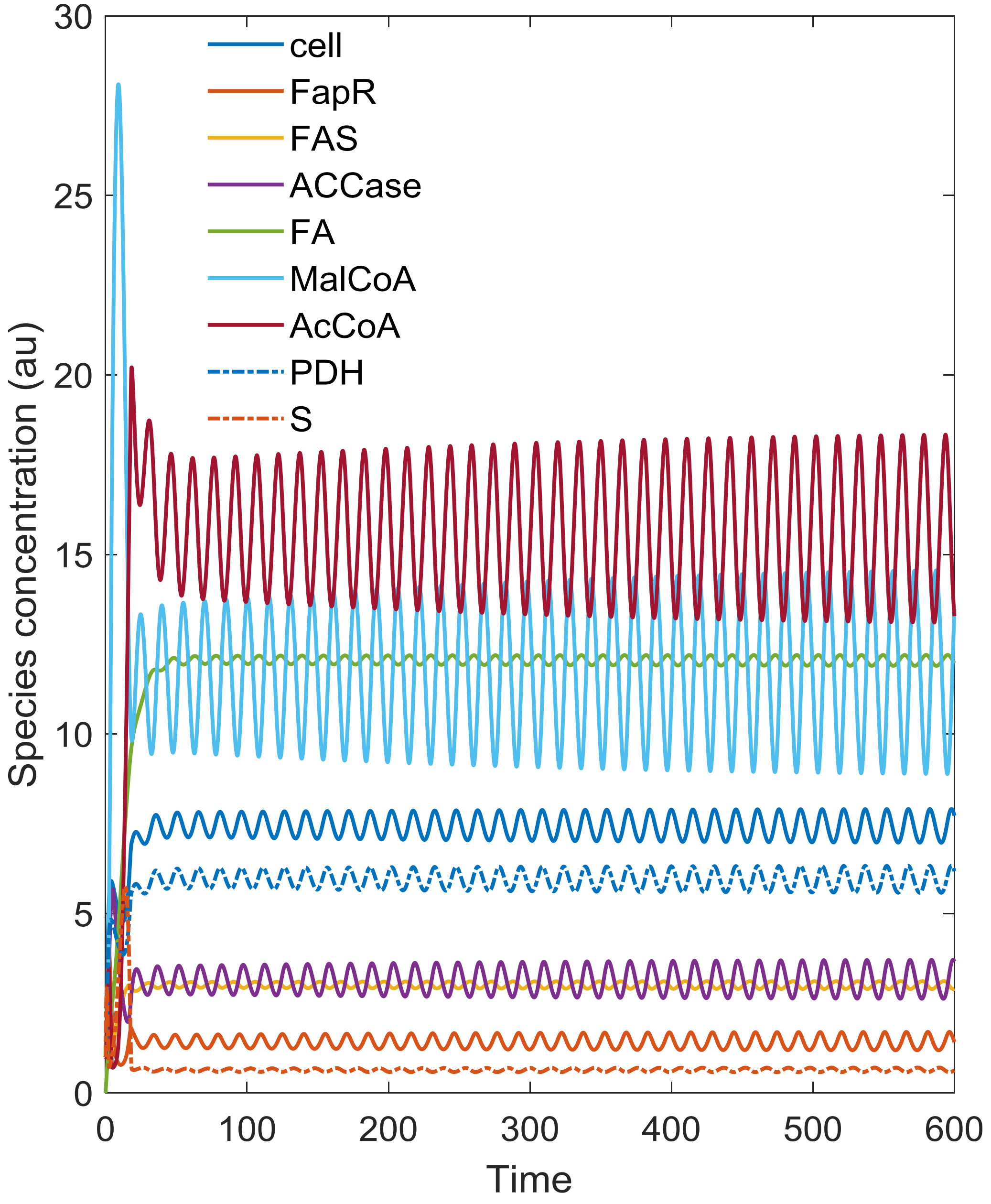


**Fig. S1**. Oscillatory pattern of species concentration for the autonomous Malonyl-CoA controller at $D$ = 0.15. Other parameters used here are the same as the parameters used in Fig. 2.

**Supplementary notes 1. Numerically evaluating the Jacobian Matrix and eigenvalues for the steady states (equilibrium points) listed in Fig. 2 and Fig. S1.**

Here is a numerical evaluation of the Jacobian matrix and the corresponding eigenvalues for the equilibrium states in Fig. 2 (*D*=0.15) and Fig. S1. All MATLAB code and output results are marked in blue color.

-----------------------

```
>> D = 0.15;
tspan = [0:0.1:600];
y1_0=1; y2_0=1; y3_0=1; y4_0=1.0; y5_0=0; y6_0=1; y7_0=1; y8_0=1;
y9_0=1;
[T,Y]= ode45(@(t,y) MalCoA_Switch(t,y,D),tspan, [y1_0 y2_0 y3_0 y4_0 y5_0...
   y6_0 y7_0 y8_0 y9_0]);
plot(T,Y(:,1),'linewidth',1.5);
hold on
plot(T,Y(:,2),'-','linewidth',1.5);
plot(T,Y(:,3),'-','linewidth',1.5);
plot(T,Y(:,4),'-','linewidth',1.5);
plot(T,Y(:,5),'-','linewidth',1.5);
plot(T,Y(:,6),'-','linewidth',1.5);
plot(T,Y(:,7),'-','linewidth',1.5);
plot(T,Y(:,8),'-.','linewidth',1.5);
plot(T,Y(:,9),'-.','linewidth',1.5);
legend('cell','FapR','FAS','ACCase','FA','MalCoA','AcCoA','PDH','S');
legend boxoff;
xlabel('Time','FontSize',16);
ylabel('Species concentration (au)','FontSize',16);
>> J1 = numeric_jacobian(@test_MalCoA_Switch,Y(5838,:))
```

% Here is the Jacobian matrices numerically computed by Matlab. The equilibrium points are approximately taken at t=583.8, t=599.0 and t = 599.1. Note that we cannot accurately retrieve the steady states due to the limitations of numerical footstep (In ODE45, we iteratively retrieve a solution every 0.1 time scale). If we could take infinitesimally small footsteps, we may approach the exact equilibrium points.

```
J1 =

   -0.0118         0         0         0         0   -0.0760         0         0    0.9786
    0.1106   -0.6367         0         0         0   -0.0672         0         0    0.7829
    0.0069   -0.2491   -0.1500         0         0   -0.0038         0         0    0.0489
    0.0069    0.9962         0   -0.1500         0   -0.0038         0         0    0.0489
         0         0    0.6000         0   -0.1500    0.0000         0         0         0
         0         0   -1.5000    2.0000         0   -0.1500    0.0000         0         0
         0         0         0   -2.0000         0         0   -0.1500    1.3818   17.5659
    0.1106         0         0         0         0   -0.0608         0   -0.1500    0.7829
   -0.2304         0         0         0         0    0.1266         0   -0.7677  -11.5399

>> J2 = numeric_jacobian(@test_MalCoA_Switch,Y(5990,:))

J2 =

   -0.0114         0         0         0         0   -0.0764         0         0    0.9826
    0.1108   -0.6365         0         0         0   -0.0677         0         0    0.7861
    0.0069   -0.2497   -0.1500         0         0   -0.0038         0         0    0.0491
    0.0069    0.9987         0   -0.1500         0   -0.0038         0         0    0.0491
         0         0    0.6000         0   -0.1500    0.0000         0         0         0
         0         0   -1.5000    2.0000         0   -0.1500    0.0000         0         0
         0         0         0   -2.0000         0         0   -0.1500    1.3805   17.6082
    0.1108         0         0         0         0   -0.0611         0   -0.1500    0.7861
   -0.2309         0         0         0         0    0.1273         0   -0.7669  -11.5700

>> J3 = numeric_jacobian(@test_MalCoA_Switch,Y(5991,:))

J3 =

   -0.0124         0         0         0         0   -0.0752         0         0    0.9701
    0.1101   -0.6370         0         0         0   -0.0664         0         0    0.7761
    0.0069   -0.2480   -0.1500         0         0   -0.0038         0         0    0.0485
    0.0069    0.9920         0   -0.1500         0   -0.0038         0         0    0.0485
         0         0    0.6000         0   -0.1500    0.0000         0         0         0
         0         0   -1.5000    2.0000         0   -0.1500    0.0000         0         0
         0         0         0   -2.0000         0         0   -0.1500    1.3863   17.4373
    0.1101         0         0         0         0   -0.0601         0   -0.1500    0.7761
   -0.2294         0         0         0         0    0.1253         0   -0.7701  -11.4542
```

% We will numerically evaluate the eigenvalues of the Jacobian Matrices at the three approximately steady states at t=583.8, t=599.0 and t = 599.1, which are equivalently to the solution Y(5838,:), Y(5990,:) and Y(5991,:).

```
>> eig1=eig(J1)

eig1 =

  -0.1500 + 0.0000i
 -11.4679 + 0.0000i
  -0.8774 + 0.0000i
  -0.0002 + 0.4121i
  -0.0002 - 0.4121i
  -0.1428 + 0.0000i
  -0.1500 + 0.0000i
  -0.1500 + 0.0000i
  -0.1500 + 0.0000i

>> eig2=eig(J2)
```

eig2 =

```
  -0.1500 + 0.0000i
 -11.4979 + 0.0000i
  -0.8787 + 0.0000i
   0.0006 + 0.4137i
   0.0006 - 0.4137i
  -0.1427 + 0.0000i
  -0.1500 + 0.0000i
  -0.1500 + 0.0000i
  -0.1500 + 0.0000i
```

```
>> eig3=eig(J3)
```

eig3 =

```
  -0.1500 + 0.0000i
 -11.3822 + 0.0000i
  -0.8751 + 0.0000i
  -0.0017 + 0.4094i
  -0.0017 - 0.4094i
  -0.1430 + 0.0000i
  -0.1500 + 0.0000i
  -0.1500 + 0.0000i
  -0.1500 + 0.0000i
```

-------------------

By evaluating the three steady states, we observed 7 of the 9 eigenvalues are negative real numbers in eig1, eig2 and eig3, except for the 4$^{th}$ and 5$^{th}$ components with imaginary parts which are highlighted in yellow. The highlighted eigenvalues for the three steady states have real parts approximating to zero with conjugate imaginary parts. For the 2$^{nd}$ steady state at t =599.0 and the 3$^{rd}$ steady state at t=599.1, we observed that the real part of the eigenvalues for J2 and J3 changed the sign. For example, the 4$^{th}$ and 5$^{th}$ components of the real part of eigenvalues changes its sign from +0.0006 in eig2 to -0.0017 in eig3, indicating that there exists a zero real part eigenvalue if we could take infinitesimally small footsteps between t=599.0 and t=599.1. Zero real part with imaginary eigenvalues indicate a stable oscillation. *Therefore, the steady state solutions are attracted to a limit cycle in the phase plane*.

**Supplementary notes 2. Plotting the steady state solutions of fatty acids, FAS and malonyl-CoA when the malonyl-CoA inhibition constant was changed.**

By plotting the steady state solutions on a 3-D phase plane, we did find looping patterns, as we change the bifurcation parameter K1 (malonyl-CoA inhibition constant). Here is the code we run,

```
for K1=0.1:0.0005:1.5;
tspan = [0:0.1:120];
y1_0=9.4338; y2_0=4.2121; y3_0=2.3945; y4_0=6.1138; y5_0=11.1925;
y6_0=11.3272; y7_0=12.2715; y8_0=7.5470;
y9_0=0.6210;

[T,Y]= ode45(@(t,y) MalCoA_Switch_K1(t,y,K1),tspan, [y1_0 y2_0 y3_0 y4_0 y5_0...
   y6_0 y7_0 y8_0 y9_0]);
plot3(Y(1200,3),Y(1200,6),Y(1200,5), 'r.');
hold on;
end
>> xlabel('FAS (au)');
>> ylabel('Malonyl-CoA (au)');
>> zlabel('Fatty acids (au)');
```

% The initial condition starts with one of the equilibrium points when K1 = 0.5. Here is the 3-D phase plane when the malonyl-CoA inhibition constant (K1) was varied from 0.1 to 1.5. This figure was also attached as a main figure in the main text (Fig. 7). The looping pattern of steady state solutions (fatty acids, fatty acid synthase and malonyl-CoA) may indicate multiplicity and hysteresis of the system when K1 was changed.

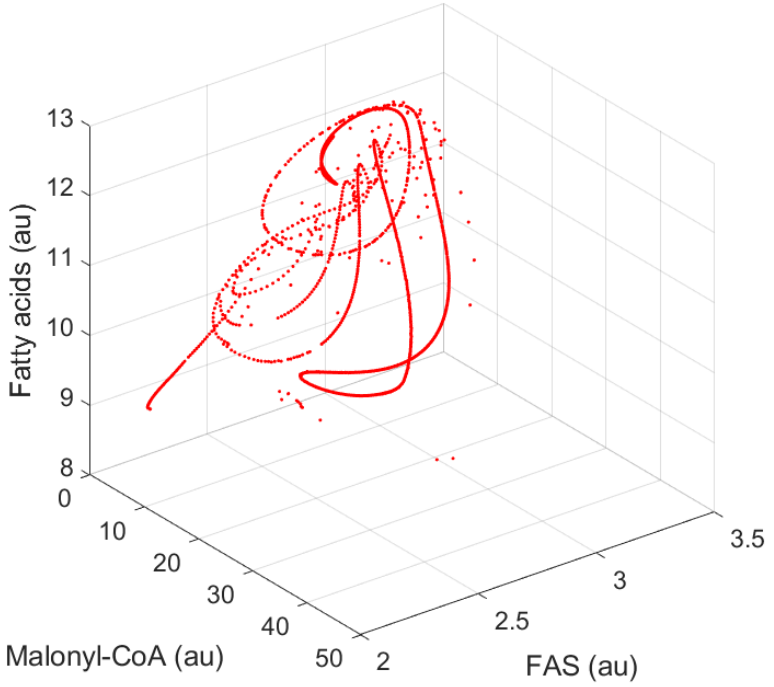

**Supplementary notes 3. Adding the trajectory with K1 = 0.3 to the 3-D phase plane.**

```
for K1=0.1:0.0005:1.5;
tspan = [0:0.1:120];
y1_0=9.4338; y2_0=4.2121; y3_0=2.3945; y4_0=6.1138; y5_0=11.1925;
y6_0=11.3272; y7_0=12.2715; y8_0=7.5470; y9_0=0.6210;
[T,Y]= ode45(@(t,y) MalCoA_Switch_K1(t,y,K1),tspan, [y1_0 y2_0 y3_0 y4_0 y5_0...
   y6_0 y7_0 y8_0 y9_0]);
plot3(Y(1200,3),Y(1200,6),Y(1200,5), 'r.');
hold on
end
xlabel('FAS (au)');
ylabel('Malonyl-CoA (au)');
zlabel('Fatty acids (au)');
hold on
% draw a specific trajectory with K1=0.3 and a different starting point
K1=0.3;
tspan = [0:0.1:120];
y1_0=1; y2_0=1; y3_0=1; y4_0=1; y5_0=1; y6_0=1; y7_0=1; y8_0=1;
y9_0=1;
[T,Y]= ode45(@(t,y) MalCoA_Switch_K1(t,y,K1),tspan, [y1_0 y2_0 y3_0 y4_0 y5_0...
   y6_0 y7_0 y8_0 y9_0]);
plot3(Y(:,3),Y(:,6),Y(:,5), 'b.');

% A specific trajectory for K1 = 0.3 was added to the above solution space, marked in blue color in the figure attached below. This figure is attached as supplementary figures Fig. S2.
```

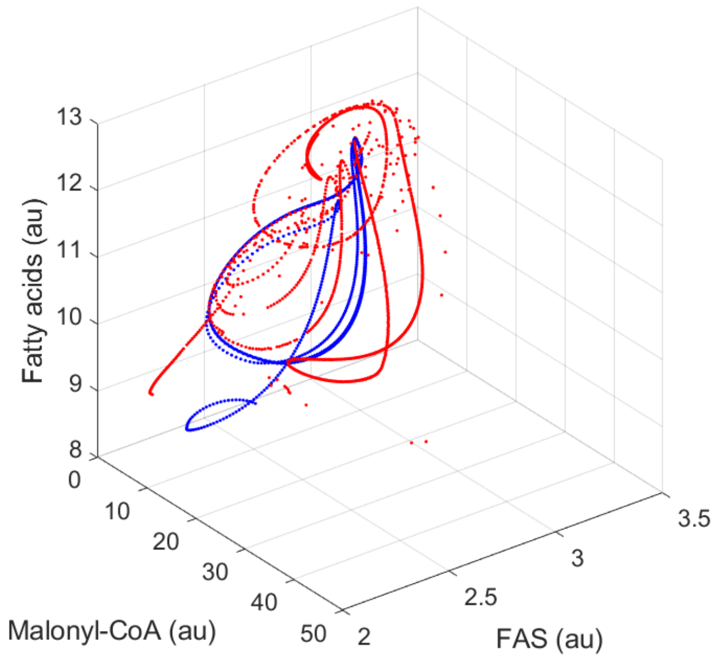
